# Ozone-induced changes in murine lung extracellular vesicle number and small RNA content

**DOI:** 10.1101/2020.06.17.157156

**Authors:** Gregory J. Smith, Adelaide Tovar, Matt Kanke, Praveen Sethupathy, Samir N. P. Kelada

## Abstract

Inhalation exposure to ozone (O_3_) causes adverse respiratory health effects that result from airway inflammation, a complex response mediated by changes to airway cellular transcriptional programs. These programs may be regulated in part by a subset of microRNAs transferred between cells (e.g. epithelial cells and macrophages) via extracellular vesicles (EV miRNA). To explore this, we exposed female C57BL/6J mice to filtered air (FA), 1, or 2 ppm O_3_ by inhalation and collected bronchoalveolar lavage fluid (BALF) 21 hours later for markers of airway inflammation, EVs, and EV miRNA. Both concentrations of O_3_ significantly increased markers of inflammation (neutrophils and total protein) and the number of EVs in the BALF. Using high-throughput small RNA sequencing, we identified several differentially expressed (DE) BALF EV miRNAs after 1 ppm (16 DE miRNAs) and 2 ppm (99 DE miRNAs) O_3_ versus FA exposure. O_3_ concentration response patterns in EV miRNA expression were apparent, particularly for the two most highly expressed (miR-2137 and miR-126-3p) and lowly expressed (miR-378-3p and miR-351-5p) miRNAs. Integrative analysis of EV miRNA expression and airway cellular mRNA expression identified EV miR-22-3p as a candidate regulator of transcriptomic responses to O_3_ in airway macrophages. In contrast, we did not identify candidate miRNA regulators of mRNA expression data from conducting airways (predominantly composed of epithelial cells). In summary, our data show that O_3_ exposure alters EV release and EV miRNA expression, suggesting that further investigation of EVs may provide insight into their effects on airway macrophage function and other mechanisms of O_3_-induced respiratory inflammation.

## Introduction

Ozone (O_3_) is a highly reactive, oxidant air pollutant associated with significant adverse respiratory health effects including airway inflammation and exacerbation of respiratory diseases such as asthma (1). Observations of the health effects of O_3_ date to the mid-19th century (47) and yet, knowledge of the underlying mechanisms is still incomplete. Given that ground level O_3_ concentrations in many regions regularly exceed the current U.S. national standard (0.07 ppm) (53), O_3_-induced airway inflammation can occur at even lower concentrations (30), and ambient O_3_ concentrations are expected to rise in the coming decades due to climate change (41), research to identify the mechanisms by which O_3_ causes respiratory health effects is imperative.

Inhaled O_3_ reacts rapidly with the airway surface lining liquid to generate reactive oxygen species, lipid peroxides, and other products that induce a complex and dynamic respiratory tract response (38, 42). Inflammation occurs up to 48 hours following exposure, initiated in part by the release of cytokines and marked by edema and the influx of neutrophils to the lung (38). Ultimately, the anti-inflammatory and pro-resolving activities of the respiratory tract and innate immune cells restore homeostasis by about 72 hours (29).

Alveolar macrophages and epithelial cells are two airway cell-types with readily apparent functional responses to O_3_ (1). Alveolar macrophages show reduced phagocytosis (17) and enhanced antigen presentation activity (32). Epithelial cell barrier dysfunction and damage are evidenced by the presence of an intra-alveolar, albumin-rich exudate following O_3_ exposure (21). Shared functional responses, albeit differing with respect to timing and precise mediators, include the release of pro-inflammatory cytokines (e.g. IL-6 (11), IL-1 family (37), TNFα (13)) and altered production/activity of other proteins (e.g. surfactant (20), metallothionein (24), matrix metalloproteinases (55)), and small molecules (e.g. antioxidants (5), eicosanoids (11), specialized pro-resolving lipid mediators (29)).

Underlying the functional cellular responses of airway macrophage and epithelial cells to O_3_ are alterations in inflammatory, immune, and oxidative stress response gene expression programs. Transcriptomic and proteomic studies have better illuminated the landscape of gene expression responses to O_3_, revealing the involvement of the protease/anti protease system (28), heat shock proteins (39), and NRF2 (8) and NF-kB family transcription factors (26). Novel O_3_-responsive genes have also been discovered recently, including oxytocin receptor (*Oxtr*) in conducting airways and hairy enhancer of split 1 (*Hes1*) in airway macrophages (51). Although the gene expression landscape of the airway response to O_3_ is becoming clearer, the precise mechanisms regulating gene expression responses to O_3_ are unknown.

Extracellular vesicles (EVs) have emerged as an important means of airway intercellular communication and may directly influence gene expression responses of the respiratory tract (18, 36, 44). These nanometer scale, lipid membrane-bound particles carry several classes of cargo including small RNAs such as microRNAs (miRNA) (10). miRNAs regulate gene expression post-transcriptionally by destabilizing or inhibiting the translation of miRNA target-mRNAs, and a single miRNA can regulate hundreds of different mRNAs (4). EV-derived miRNAs (EV miRNAs) are readily detectable in the airway lumen and are taken up by respiratory tract cells including airway macrophages and epithelial cells (18). As such, EVs represent a unique mechanism for the transfer of miRNA, and intercellular miRNA-mediated regulation of airway gene expression.

Inflammatory lung diseases including asthma (35), idiopathic pulmonary fibrosis (40), and chronic obstructive pulmonary disease (COPD)(16) as well as oxidant- or allergen-induced lung inflammation (33, 43) are associated with altered airway EV miRNA profiles. In addition, there is growing evidence that EV miRNAs directly elicit airway cellular gene expression and phenotypic effects. For example, studies have demonstrated their direct effects on myofibroblast and macrophage differentiation (25, 46) and protein secretion by epithelial cells (19). Thus, it is likely that EV miRNA play functional roles in O_3_-induced inflammation (2); however, the extent to which O_3_ exposure affects EV miRNA expression is unknown. We hypothesized that O_3_ exposure dysregulates airway EV miRNAs and that specific dysregulated EV miRNAs correlate with their target mRNA expression changes in airway cells. To investigate this possibility, we conducted high-throughput small RNA sequencing of miRNA isolated from murine airway EVs following filtered air (FA), 1 and 2 ppm O_3_ exposure and used an integrative bioinformatics approach to identify putative EV miRNA regulators of gene expression in airway macrophages and conducting airways. Here, we provide both an important initial quantification of the murine airway EV miRNA landscape and identify several EV miRNAs as putative regulators of O_3_-induced gene expression changes.

## Materials and Methods Animals

Female C57BL/6J mice at 8 to 10 weeks of age were purchased from the Jackson Laboratory (Bar Harbor, Maine) and used for all experiments. Mice were housed in cages at a maximum density of 5/cage on ALPHA-Dri bedding (Shepard) under standard 12h lighting conditions, with *ad libitum* access to food (Envigo 2929) and water. After a 1-week acclimation period, mice were randomly assigned to different exposure groups. Whole body inhalation exposures to filtered air (FA), 1 or 2 ppm O_3_ were conducted as previously described (48, 51).

Environmental conditions for each experiment, including mean O_3_ concentration (Supplemental Figure 1), temperature, and relative humidity were recorded (Supplemental Table 1). For tissue collection and phenotyping, mice were euthanized with a lethal dose of urethane (2 g/kg, i.p.) followed by exsanguination. The sample size for each group is indicated in the figure legends. All experiments were conducted within an AAALAC approved facility and were approved by the Institutional Animal Care and Use Committee (IACUC) at the University of North Carolina at Chapel Hill.

### Bronchoalveolar lavage

Following euthanasia, a 21g catheter was inserted in the tracheal lumen 2-cartilage rings posterior to the larynx, towards the lung of the mice. The lung was then gently lavaged twice (1 x 0.5mL, 1 x 1mL) with ice-cold PBS containing cOmplete protease inhibitor cocktail (Roche) using a 1-mL syringe. Collected bronchoalveolar lavage fluid (BALF) was centrifuged (400 x g, 10 mins, 4°C) to pellet cells. Red blood cells were lysed with hypertonic saline and the remaining white blood cells were resuspended in Hank’s Balanced Salt Solution and counted by hemocytometer and mounted on slides via cytospin. Cytospin slides were stained with Kwik-Diff (Shandon). Two investigators, blinded to exposure group, performed differential cell counts (minimum 300 cells per slide).

### EV isolation and characterization

After the removal of cells (above), a second centrifugation of the BALF was performed (16,000 x g, 10mins, 4°C) to pellet cellular debris and larger vesicles (e.g. apoptotic bodies). The concentration and size distribution of particles within the BALF supernatants were then measured prior to EV isolation. Nanoparticle tracking analysis (NTA) was performed on a Nanosight NS500 (Malvern Panalytical) at the UNC Nanomedicines Characterization Core Facility. BALF supernatants were diluted 1:100 in 0.02-μM filtered PBS prior to loading on the NTA instrument. NTA data were derived from five separate, 40-second motion captures using Nanosight software v3.2 (Malvern Panalytical).

### EV small RNA isolation and expression analysis

BALF samples were pooled into groups of three to four and total RNA was isolated from pooled samples using an ExoRNeasy Serum/Plasma Midi kit (Qiagen). The ExoRNeasy kit purifies EVs from biofluids using affinity binding columns and total RNA from lysed EVs using an RNA binding column (12). The yield and integrity of small RNA from isolated and lysed EVs were measured using a small RNA chip on an Agilent 2100 Bioanalyzer prior to RNA sequencing, or by Nanodrop 2000 and Qubit high sensitivity RNA assay (Thermofisher) prior to use in RT-qPCR assays. Small RNA sequencing libraries were prepared using a BioO Scientific NEXT Flex-v3 kit. Single end sequencing was performed on the Illumina HiSeq 4000 platform at the UNC High-Throughput Sequencing Facility (University of North Carolina, Chapel Hill, NC, USA). Using miRquant 2.0 (27), small RNA reads were trimmed of adapters, aligned to the *Mus musculus domesticus* reference genome (assembly NCBI37/mm9), and miRNAs and isomiRs (miRNA sequence variants) were quantified. Small RNA Bioanalyzer results, read length and mapping statistics are presented in the results section and Supplemental Figures 2 and 3.

For RT-qPCR, cDNA synthesis was performed on 10 ng of RNA using a TaqMan advanced miRNA cDNA Synthesis kit (ThermoFisher Scientific). miRNA qPCR reactions were performed in triplicate on a BioRad CFX 384 Touch Real-Time PCR detection system using TaqMan Fast Advanced qPCR Master Mix (ThermoFisher Scientific) according to the manufacturer’s instructions. TaqMan Advanced miRNA assays for miR-22-3p (mmu481004_mir) and miR-23a-3p (mmu478532_mir) were used for RT-qPCR expression analysis. miR-23a-3p was used for normalization based on its low coefficient of variation in expression across samples measured by small RNA sequencing. miRNA levels are expressed as relative quantitative values (RQVs) and statistical analyses were conducted using dCq values.

### EV miRNA target mRNA prediction

miRNA target site enrichment analysis was performed on lists of O_3_-responsive genes identified in our previously published work (51) using the bioinformatics algorithm mirRhub (3). Using, for example, a list of down regulated genes as an input, miRhub determines if any miRNAs are significantly over represented in the 3’ UTR TargetScan-predicted miRNA binding sites of the genes as compared to randomly generated gene lists. Identified miRNAs would be expected to be differentially expressed in the opposite direction of the gene list input.

## Statistical Analysis

For analysis presented in Supplemental Figure 2 and Figure 1, raw data were subjected to Box-Cox transformation prior to statistical testing using ANOVA and pairwise t-tests in R (version 3.6.1). The results of a test were considered significant if the *p*-value was < 0.05.

## Results

### Ozone (O_3_) inhalation causes inflammation and increased extracellular vesicle (EV)-sized particles in the lung

We exposed female C57BL6/J mice to filtered air (FA), 1 or 2 ppm O_3_ for 3h and collected bronchoalveolar lavage fluid (BALF) 21h later for isolation of EVs and measurement of lung inflammation and injury. BALF samples from both 1 and 2 ppm O_3_ exposed mice exhibited characteristic pulmonary inflammatory phenotypes including significant increases in cellularity, driven by macrophages and neutrophils, and injury as measured by increased total protein compared to FA (control) mice (Supplemental Figure 2). As expected, both tissue injury and inflammation were most severe in the 2 ppm O_3_ exposed group, compared to the 1 ppm O_3_ or FA groups.

We measured the number of particles in BALF across a size range of ∼50-1000 nm by nanoparticle tracking analysis (NTA). We observed a concentration-dependent increase in the total number of BALF particles in the size range of roughly 50-400 nm (Figure 1A). The increase in BALF particle concentration was statistically significant at 2 ppm O_3_ compared to the 1 ppm O_3_ and FA groups (Figure 1B). Additionally, we observed modest shifts in the particle size distributions across treatment groups, with both the means and modes of particle size decreasing from FA to 2 ppm O_3_ (Figures 1C and D).

**Figure 1.**
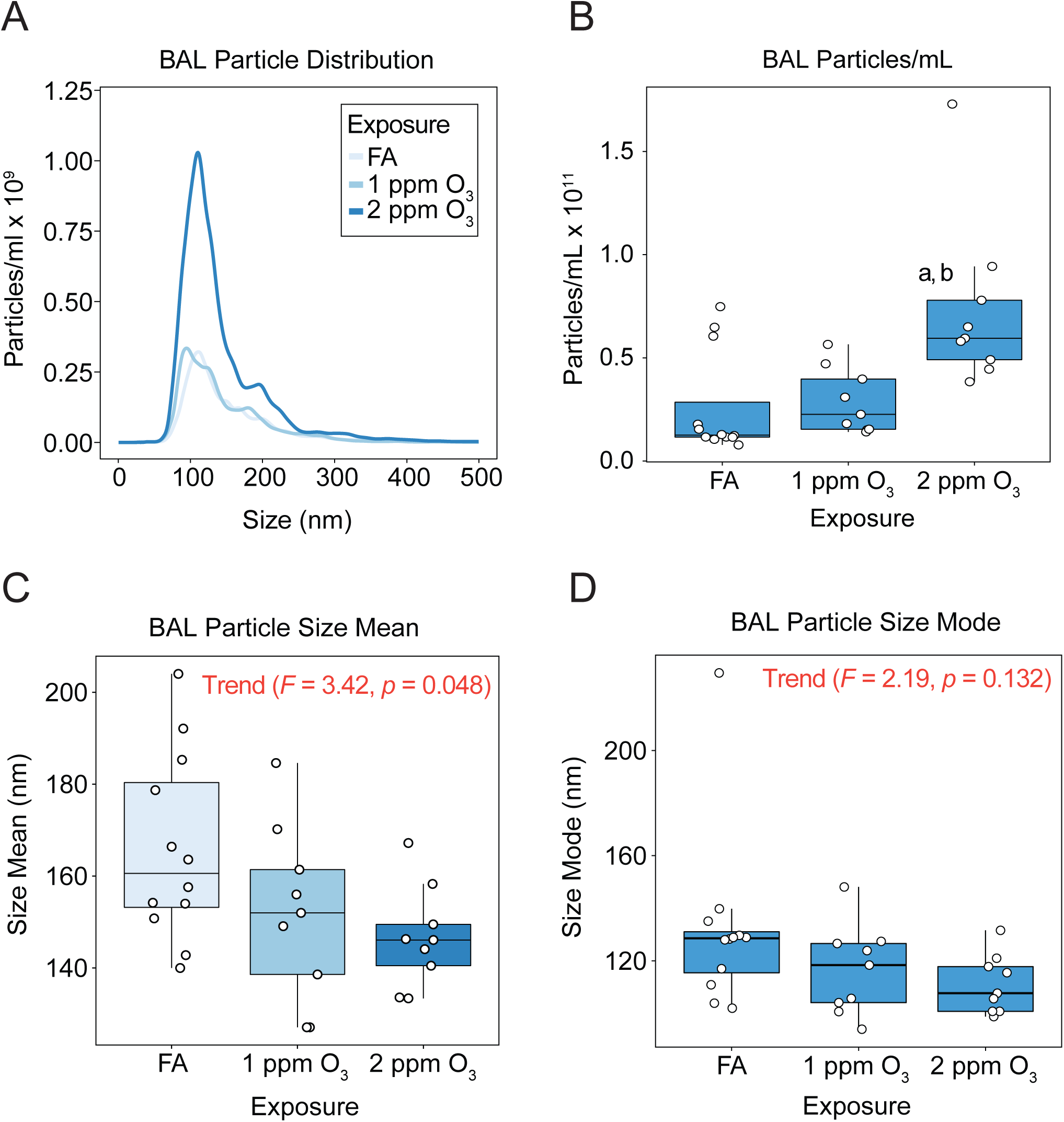
O_3_ exposure increases in extracellular vesicle-sized particles in the lung. Nine to ten-week-old female C57BL/6J mice were exposed to filtered air (FA; n = 12), 1 (n = 9) or 2 (n = 9) ppm O_3_ for 3h and bronchoalveolar lavage fluid was collected at 21h for isolation and analysis of extracellular vesicles. (A) Size distribution, (B) total concentration, (C) size mean, and (D) size mode of particles in BALF samples were measured by nanoparticle tracking analysis. Results are depicted in (A) as the mean particle-size distribution. In (B-D) box-and-whisker plots depict the minimum, first quartile, median, third quartile, and maximum of the data with all points overlaid. a: *p* < 0.05 compared to FA group, b: *p* < 0.05 compared to 1ppm group.

### Ozone (O_3_) induces miRNA expression changes in lung Evs

To obtain sufficient quantities of small RNA for downstream sequencing, we pooled BALF from three to four mice per treatment group, isolated EV-total RNA from each of the pools, and characterized RNA abundance and size distributions. RNA abundance did not differ between groups (Supplemental Figure 3A), and RNA contents from each pool were predominantly in the size range of 20-30 nucleotides, the size of mature miRNA (Supplemental Figure 3B). To identify O_3_-responsive EV miRNAs, we performed high-throughput small RNA sequencing (small RNA-seq) on each of the pooled EV RNA samples (n=3 pools/exposure group). Small RNA-seq yielded an average of 44 million reads per sample and, after trimming and exclusion of reads outside 14-40 nucleotides, an average of 78% of reads were mapped across exposure groups (Supplemental Figure 4A). Read statistics indicate that 1-2% of reads mapped to miRNA, while the bulk of reads (∼70-90%) were in the range of 30-35 nucleotides and mapped to tRNA, suggestive of tRNA-derived RNA fragments (tDRs) (Supplemental Figures 4B and C). The balance of reads consisted of other small, unannotated RNAs (e.g., yRNAs). Across exposure groups, we observed an increase in the percentage of reads mapped to these other small RNAs and a decrease in reads mapped to tRNAs, while the percentage of miRNA was reduced by half in the 2 ppm O_3_ exposure group from ∼2 to ∼1%.

We performed principal components analysis (PCA) on the 50 most variably expressed EV miRNAs, which revealed distinct EV miRNA expression patterns by exposure (Figure 2A). Using fold change of ± 2, adjusted *p* value < 0.05, and an expression level of >50 reads per million miRNAs mapped (RPMMM) as thresholds, we identified 16 significantly differentially expressed EV miRNAs after 1 ppm O_3_ exposure and 99 significantly differentially expressed EV miRNAs after 2 ppm O_3_ exposure. Overall, we observed a greater proportion of downregulated versus upregulated EV miRNAs in both the 1 and 2 ppm O_3_ exposed groups compared to FA (Figures 2B and C). Hierarchical clustering of the 50 most variably expressed EV miRNAs revealed several different concentration-response patterns of expression including monotonic and non-monotonic relationships (Figure 3A), as we previously observed for mRNAs (51). A threshold effect was observed for the bulk of EV miRNAs, illustrated by the closer correlation of the FA and 1 ppm O_3_ groups and greater level of dysregulation in the 2 ppm O_3_ group (Figure 3A). The most highly upregulated EV miRNA after both 1 and 2 ppm O_3_ exposure was miR-2137 (Figure 3B) as well as several miR-2137 isomiRs generated by alternative processing or nucleotide additions (Supplemental Figure 5). In addition to miR-2137, miR-126-3p was one of the few upregulated miRNAs overall (∼3.0 and 4.4 fold, 1 and 2 ppm O_3_ versus FA) (Figure 3B). The most highly downregulated miRNAs were miR-378-3p (∼6.8 and 2.3 fold, 1 and 2 ppm O_3_ versus FA) and miR-351-5p (∼1.5 and 5.3 fold, 1 and 2 ppm O_3_ versus FA) (Figure 3B).

**Figure 2.**
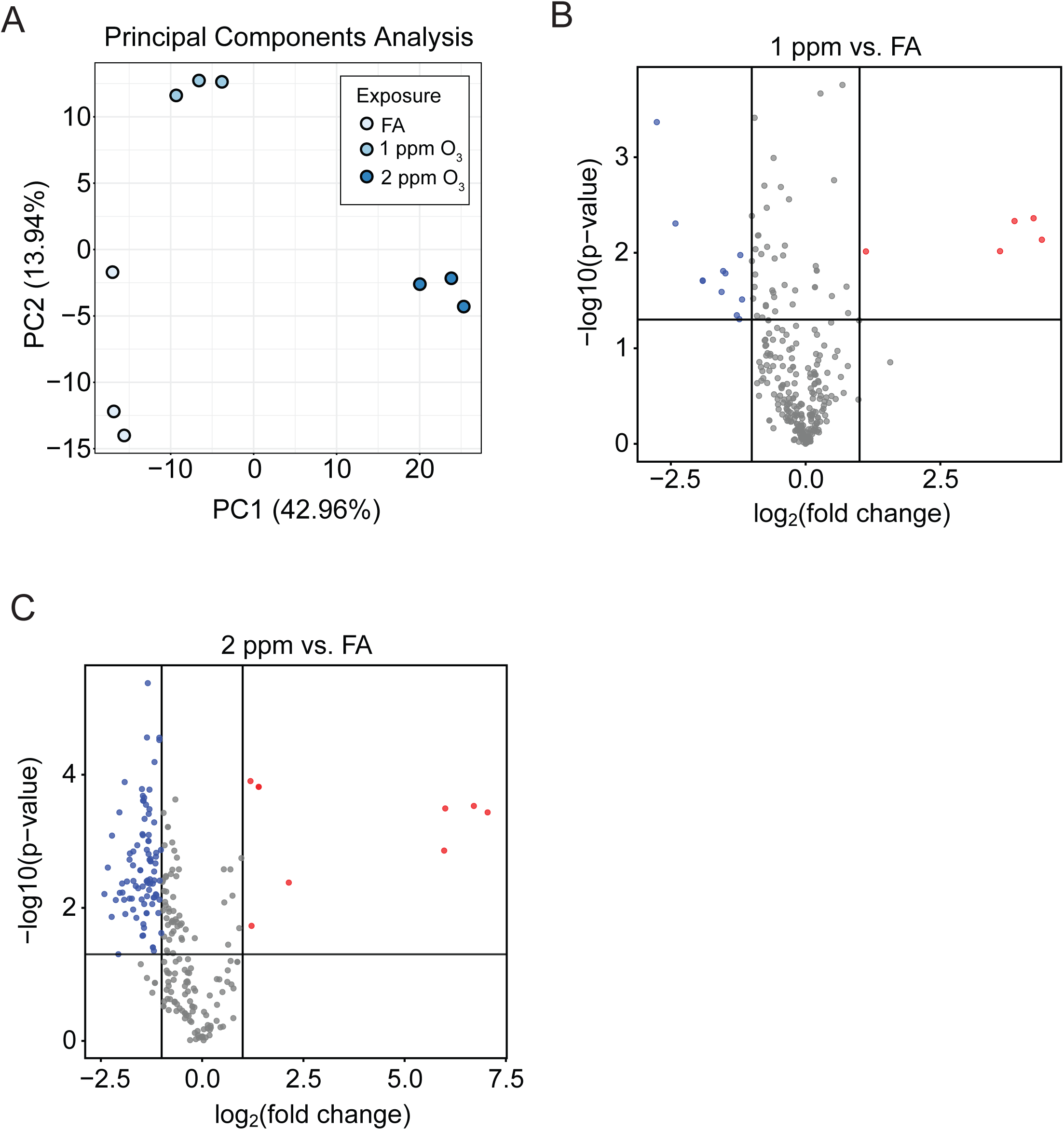
O_3_ exposure alters airway extracellular vesicle miRNA expression. (A) Principal components analysis showing separation of pooled airway EV samples by exposure group. (B and C) Volcano plots showing differentially expressed (DE) miRNAs in 1 ppm O_3_ versus FA (16 DE miRNAs) and 2 ppm O_3_ versus FA (99 DE miRNAs), respectively (horizontal line: p = 0.05, vertical lines: fold change of ± 2). Three pooled and isolated EV-RNA samples were analyzed per exposure group.

**Figure 3.**
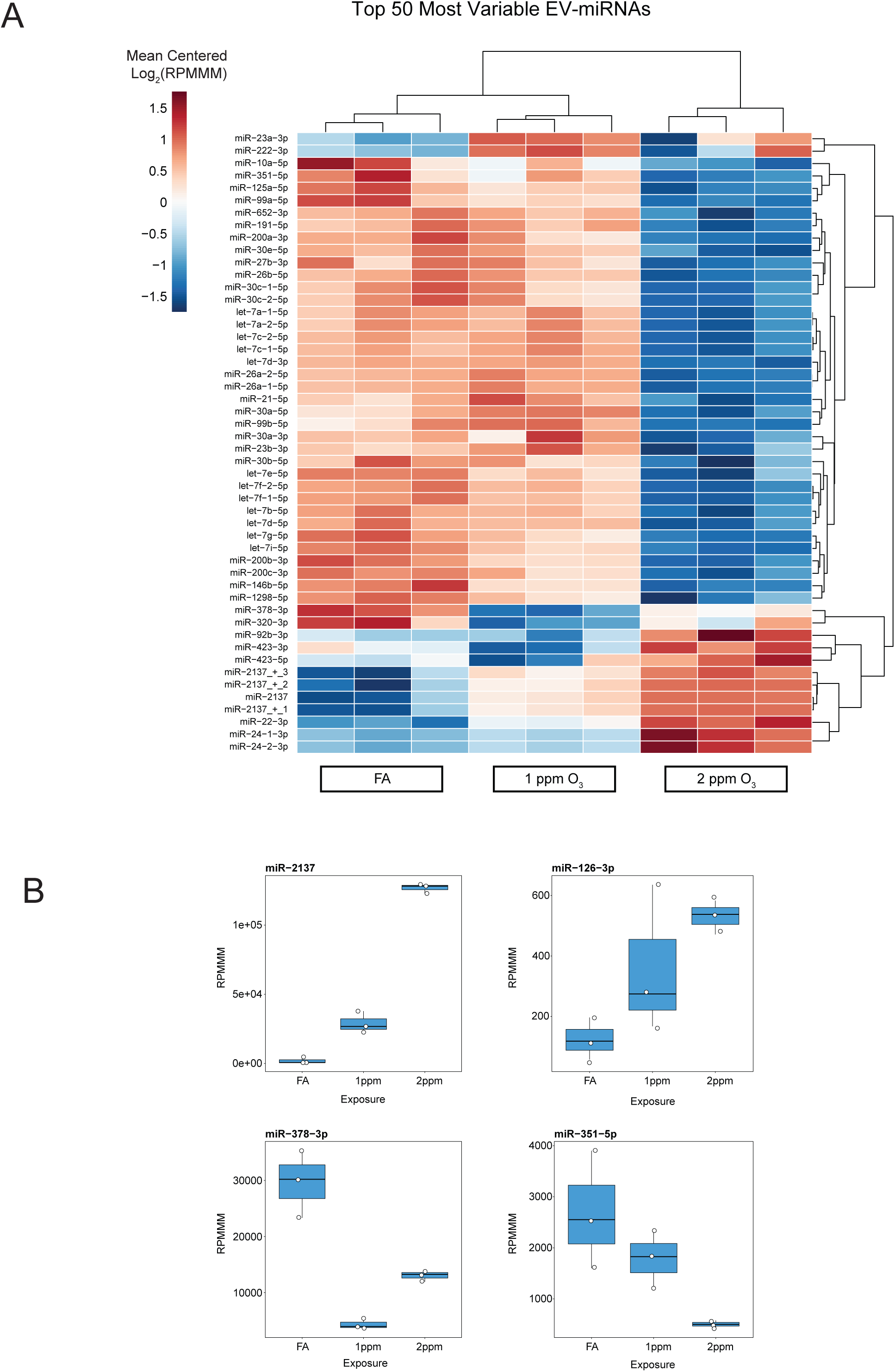
Specific O_3_-induced alterations of the EV-miRNA landscape. (A) Unsupervised hierarchical clustering analysis of the top 50 most variable EV-miRNAs. Data are presented as mean-centered, log_2_-transformed reads per million miRNAs mapped (RPMMM) for each miRNA. (B) Expression of selected, O_3_-responsive EV-miRNAs. Box-and-whisker plots depict RPMMM summarized by the minimum, first quartile, median, third quartile, and maximum with all points overlaid. n = 3 pooled samples per group.

### Integration of EV miRNA and tissue mRNA data to identify putative regulatory networks

Previously, we characterized the transcriptional profiles of airway macrophages and conducting airways of mice exposed to O_3_ (51). We integrated those mRNA expression data with the EV miRNA data presented here as a strategy to identify EV miRNAs that may cause the observed changes in tissue mRNA expression. Specifically, we sought to identify EV miRNAs that could plausibly regulate sets of target mRNAs in recipient airway macrophage or conducting airway tissue. Given the canonical role of miRNAs in decreasing mRNA levels and/or inhibiting mRNA translation (4), we focused our analysis on EV miRNAs whose expression was directionally opposed to the tissue mRNA patterns across the exposure groups (i.e., we paired miRNAs that increased after 1 and 2 ppm O_3_ with mRNAs that correspondingly decreased, or vice versa), and we stipulated that both miRNA and mRNA must be significantly differentially expressed for the designated contrast.

Our miRhub analysis identified several miRNAs that were predicted to target sets of genes in airway cells (the full set of results are shown in Supplemental Figure 6). miR-22-3p emerged as a particularly interesting candidate regulator of gene expression in airway macrophages (Figure 4) based its target enrichment score, and its relatively high expression level (>1,000 RPMMM). The pattern of O_3_-induced expression of EV miRs-22-3p is shown alongside one of its predicted target genes, *Arhgap26*, identified by miRhub (Figure 4B). Of note, in a follow-up experiment in which we exposed mice to FA or 2 ppm O_3_, we replicated our finding that EV miR-22-3p was upregulated after 2 ppm O_3_ exposure (Supplemental Figure 7).

**Figure 4.**
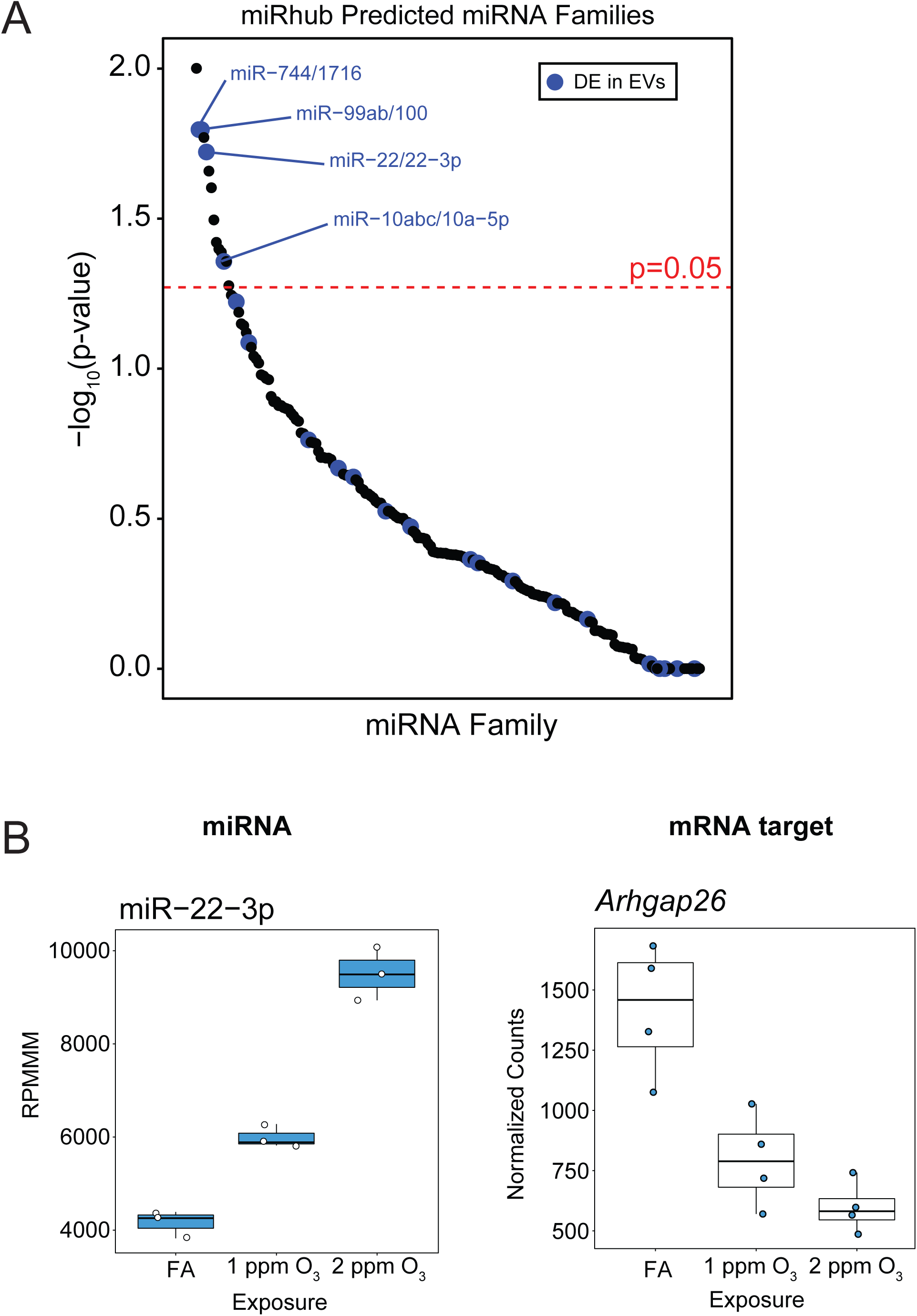
Identification of candidate EV-miRNA regulators of O_3_-induced transcriptional responses. (A) Representative plot showing output of the miRhub algorithm used to identify miR-22-3p as a potential candidate regulatory hub from a list of downregulated genes (2 ppm O_3_ versus FA) genes in airway macrophages following O_3_ exposure. All lists included genes with a fold change of ± 2, and *p* value < 0.05. Data points are plotted as –log_10_(*p*-value) generated from miRhub by miRNA family, and are highlighted if a member of the miRNA family was found to be differentially expressed in airway EVs. (B) Plot depicting RPMMM for EV miR-22-3p alongside predicted airway macrophage mRNA target expression (normalized counts from Deseq2). n=3 or 4 (mRNA data) pooled samples per group.

## Discussion

We characterized the extracellular vesicle (EV) profile and EV small RNA content following exposure to filtered air (FA), 1 or 2 ppm O_3_. Our results demonstrate a clear increase in the number of EVs in the lungs of mice following O_3_ exposure in parallel with markers of inflammation (macrophages, neutrophils, and total protein), indicating that EVs are O_3_-responsive and suggesting that they may play a role in regulating O_3_-induced inflammation.

Collectively, EVs are a heterogeneous population including two known vesicle subtypes, exosomes and microvesicles (44). In general, exosomes are smaller, around 40-100 nm in diameter, while microvesicles can be up to 1,000 nm. Size has been used to specify the subtype of EV being examined in several studies involving airway EVs (31, 34, 50). In our experiments, O_3_ exposure caused a decrease in the EV size mode from ∼120 to 100 nm, suggesting that effects of O_3_ exposure were primarily on exosomes. However, we chose to profile small RNA from the broader EV population for two main reasons: first, the mouse BALF EV small RNA profile has not been examined using high throughput sequencing methods and second, methods for isolating EV subtypes are still in a period of rapid advancement. Previous studies that focus on isolated EVs, even those using ‘gold-standard’ ultracentrifugation techniques, vary greatly and are difficult to compare directly (6, 12). We used a straightforward and commercially available isolation method that excludes non-EV RNA protein aggregates (unlike ultracentrifugation), which has yet to be described in use with mouse BALF samples. Moreover, by isolating a less biased population of EVs, we were able to observe global changes in airway luminal EV miRNA expression. We believe both considerations will help facilitate comparison and reproduction of our work.

Using small RNA-seq, we identified a host of differentially expressed (DE) airway EV miRNAs following O_3_ exposure. The overall number of DE EV miRNAs and magnitude of change in their expression was concentration-dependent. The majority of DE miRNA were downregulated, which may reflect either a selective decrease in EV loading or a loss of the miRNA’s cellular source due to toxicity. miR-2137 was identified as the most highly upregulated and expressed EV miRNA across both the 1 (∼14 fold) and 2 ppm O_3_ (∼120 fold) groups. Recently, differential expression of miR-2137 was observed in bone marrow-derived macrophages after infection with *Porphyromonas Gingivalis* bacteria (23), and inhibiting miR-2137 increased expression of the anti-inflammatory cytokine IL-10 suggesting that miR-2137 may be pro-inflammatory following O_3_ exposure. We also observed an O_3_-induced upregulation of EV miR-126-3p, a well-studied pro-inflammatory miRNA (15). miR-126-3p is also upregulated in inflammatory bowel disease (IBD), in which it targets nuclear factor-kappaB inhibitor alpha (IκB) (14, 54) and impairs intestinal epithelial barrier function (7). We observed a downregulation of miR-378-3p, whose overexpression has been shown to negatively regulate macrophage proliferation (45). Therefore, the known O_3_-induced increase in macrophage proliferation is consistent with our results and a potential regulatory effect of EV miR-378-3p (22). miR-351 was also downregulated, and recent evidence suggests that downregulation of miR-351-5p is anti-inflammatory in ischemia/reperfusion injury (57). As such, miR-351-5p may represent a pro-resolving signal after O_3_ exposure. In general, our data suggest that lung EVs collected 21 hours following O_3_ exposure contain a mixture of both pro-inflammatory and homeostatic signals.

Our integrative bioinformatics analysis of EV miRNA expression identified miR-22-3p as a putative regulator of airway macrophage gene expression following O_3_ exposure. A target of miR-22-3p, *Arhgap26*, exhibited a pattern of expression consistent with miRNA regulation. Interestingly, miR-22-3p is differentially expressed across the continuum of polarized human macrophages (9), suggesting that it may regulate macrophage polarization during O_3_ exposure. Surprisingly, although miRhub predicted some miRNA regulators for mRNAs in conducting airway tissue, none of these miRNAs were differentially expressed in EVs due to O_3_ exposure. This could be due to timing; for example, an upregulated EV miRNA could have influenced gene expression in conducting airway tissue at a different time point and is no longer or not yet differentially expressed in EVs at 21 hours. This may explain why miR-2137 was not a predicted regulatory hub in either tissue despite its high expression in EVs. Although a number of different cell types (e.g. ciliated epithelium, goblet cells, club cells, etc.) are represented in conducting airway tissue samples, the power to detect miRNA-mRNA target relationships may be reduced by obfuscation or exclusion of critical cell populations. It is also plausible that, on average, EV miRNA communication may have a greater effect on macrophages at this time point than cells of the conducting airways.

In addition to specific miRNA expression, deep sequencing also provided data on the proportion of small RNA species in BALF EVs by type: miRNA, tRNA, or other (e.g. piRNA, yRNA, etc.). We found the majority of BALF EV small RNA reads, from 60-90% across exposure groups, mapped to tRNAs. A recent study reported a high percentage of tRNA-mapped reads and EV miRNA content on the order of 10-20% in rodent serum EVs (56). As we are the first to report small RNA-seq data for murine BALF EV samples, the relatively minimal miRNA content (1-2%) we observed compared to previous serum EV small RNAseq studies was unexpected. Interestingly, the tRNA content of our samples is consistent with tRNA derived RNA fragments (tDRs) based on read length. The biology of tDRs is not well understood; however, evidence suggests they may function similarly to miRNAs by regulating gene expression through RNA interference among other suggested functions (49). The precise annotation and quantification of O_3_-induced murine EV tDRs is likely to reveal additional biological insights, particularly if specific tDRs are differentially expressed in EVs during airway inflammation.

EV small RNA content varies by the type of sample, whether collected *in vitro* (e.g. cell type) or *in vivo* (e.g. blood, urine, saliva, cell culture supernatant, etc.) (52). Therefore, it is likely that in the lung, EV small RNA content also varies by origin cell type and/or anatomical region. Because we examined a broad population of airway EVs collected from BALF, we could not characterize specific regional or cell-type differences in EV small RNA proportions. Future studies should delineate the regional and cellular sources of EVs using flow-cytometry and fluorescence microscopy-based approaches. These techniques coupled with microdissection would allow for investigation of regional differences in airway EV communication. Nevertheless, with our global EV small RNA-seq approach, we identified specific miRNAs that may regulate transcriptional responses to O_3_.

In conclusion, we show that the release of airway EVs and dysregulation of EV miRNA content are features of respiratory tract response to O3 exposure. In addition to O3-induced increases in the number of EVs and EV miRNA content, O3 also induced broad changes in the quantity of other EV small RNA species such as tRNAs. Our preliminary, hypothesis-generating work adds to the growing body of evidence that EV miRNA are both altered by and regulate inflammation in a range of mucosal tissues, including airway responses to O3 and provide compelling support for a range of mechanistic follow-up studies. In particular, addressing the question of whether changes in EV number and/or content, including miR-22-3p specifically, are responsible for changes in macrophage gene expression and the degree of O3-induced airway inflammation.

## Acknowledgements

The authors acknowledge the expert assistance of Kathryn McFadden and Courtney Nesline for assistance with mouse experiments, Dr. Rowan Beck for assistance with miRhub analysis, Dr. Mike Love for consultation on analysis of RNA-seq data, and the UNC High-Throughput Sequencing Facility (library preparation and small RNA-seq).

## Author Contributions

G.J.S. conceived and designed studies with guidance from S.N.P.K and P.S.; G.J.S. and A.T. performed experiments; G.J.S and M.K. analyzed data; G.J.S prepared the manuscript; all authors approved the final version of the manuscript.

