## Supplemental Material for "Ozone-induced changes in murine lung extracellular vesicle number and small RNA content"

**Supplemental Table 1.** Exposure Characteristics

| Exposure | Ozone (ppm) | | Relative Humidity (%) | | Temperature (C°) | |
|---|---|---|---|---|---|---|
| | *Mean* | *SD* | *Mean* | *SD* | *Mean* | *SD* |
| FA-1 | < 0.01 | - | 42.57 | 1.10 | 24.93 | 0.36 |
| FA-2 | < 0.01 | - | 42.50 | 1.26 | 25.18 | 0.36 |
| 1 ppm O3 | 0.997 | 0.015 | 43.27 | 1.06 | 24.77 | 0.23 |
| 2 ppm O3 | 1.992 | 0.020 | 42.92 | 0.91 | 24.95 | 0.25 |

Data are presented as arithmetic mean and standard deviation (SD) calculated from the time the concentration of $O_3$ reached 90% of nominal ($T_{90}$) until generation was stopped ($T_{Off}$).

### Supplemental Figure Legends

**Supplemental Figure 1.** ***$O_3$ exposure concentration-time profiles.*** Data are presented as 30-second running averages of the $O_3$ concentration (ppm) for both the (A) 1 ppm and (B) 2 ppm $O_3$ exposures. Vertical dashed lines represent the times $O_3$ reached 90% of nominal ($T_{90}$) and generation was stopped ($T_{Off}$) and horizontal dashed lines represent ± 3% of the nominal exposure concentration.

**Supplemental Figure 2.** ***$O_3$ exposure causes airway inflammation and injury.*** Nine to ten-week-old female C57BL/6J mice were exposed to filtered air (FA; n = 12), 1 (n = 9) or 2 (n = 9) ppm $O_3$ for 3 hours and bronchoalveolar lavage fluid (BALF) was collected at 21 hours for differential cell counting and total protein analysis. (A) Total number of cells, (B) percent neutrophils, (C) number of neutrophils, and (D) total protein in BALF samples. a: $p < 0.05$ compared to FA group, b: $p < 0.05$ compared to 1ppm group.

**Supplemental Figure 3.** ***Airway EV small RNA characteristics.*** Total RNA was isolated from pooled airway EV samples (n = 3) from mice exposed to filtered air (FA; n = 12), 1 (n = 9) or 2 (n = 9) and analyzed using an Agilent Bioanalyzer Small RNA Chip. (A) Plot depicting small RNA concentration data for each set of pooled EV samples. (B) A representative electropherogram from one pooled sample. Tags represent areas under the curve calculated between the vertical dashed lines for each peak. Sample peak areas are compared to that of a lower marker (green peak label) to calculate concentrations for each sample.

**Supplemental Figure 4.** ***EV Small RNA Read Statistics.*** EV small RNA sequencing reads were trimmed of adapters, and aligned to the *Mus musculus domesticus* reference genome (assemble NCBI37/mm9) using the miRquant 2.0 pipeline. (A) Plot depicting the percent of small RNA-seq reads at each step of the analysis pipeline. (B) Plot depicting the mean read lengths for all small RNAs mapped to the genome. (C) Pie-charts indicating the percentage of each small RNA species by exposure group.

**Supplemental Figure 5.** ***$O_3$-induced EV miR-2137 isoMir expression.*** Data are depicted as $log_{10}$(RPMMM) for the mature miR-2137 (solid border) and its isoMirs (dashed border) across each exposure group.

**Supplemental Figure 6.** ***miRhub analysis output.*** Plots depict the output of the miRhub algorithm for all lists used (one plot per list). Lists included all differentially expressed genes (All DE), downregulated genes (Down), upregulated genes (Up), or trending genes (Downtrend or Uptrend) for airway macrophages (solid border) or conducting airway tissue (dashed border). Lists included genes with a fold change of ± 2, and $p$ value < 0.05. Data points are plotted as $-log_{10}$($p$-value) generated from the miRhub by miRNA family, and points are highlighted if a member of the miRNA family was found to be differentially expressed in airway EVs.

**Supplemental Figure 7.** ***qRT-PCR validation of miR-22-3p expression.*** Taqman assays were used to determine the expression of miR-22-3p relative to a normalization miRNA, miR-23a-3p, in EVs from mice exposed to 2 ppm $O_3$ versus FA.

**Supplemental Figure 1.**

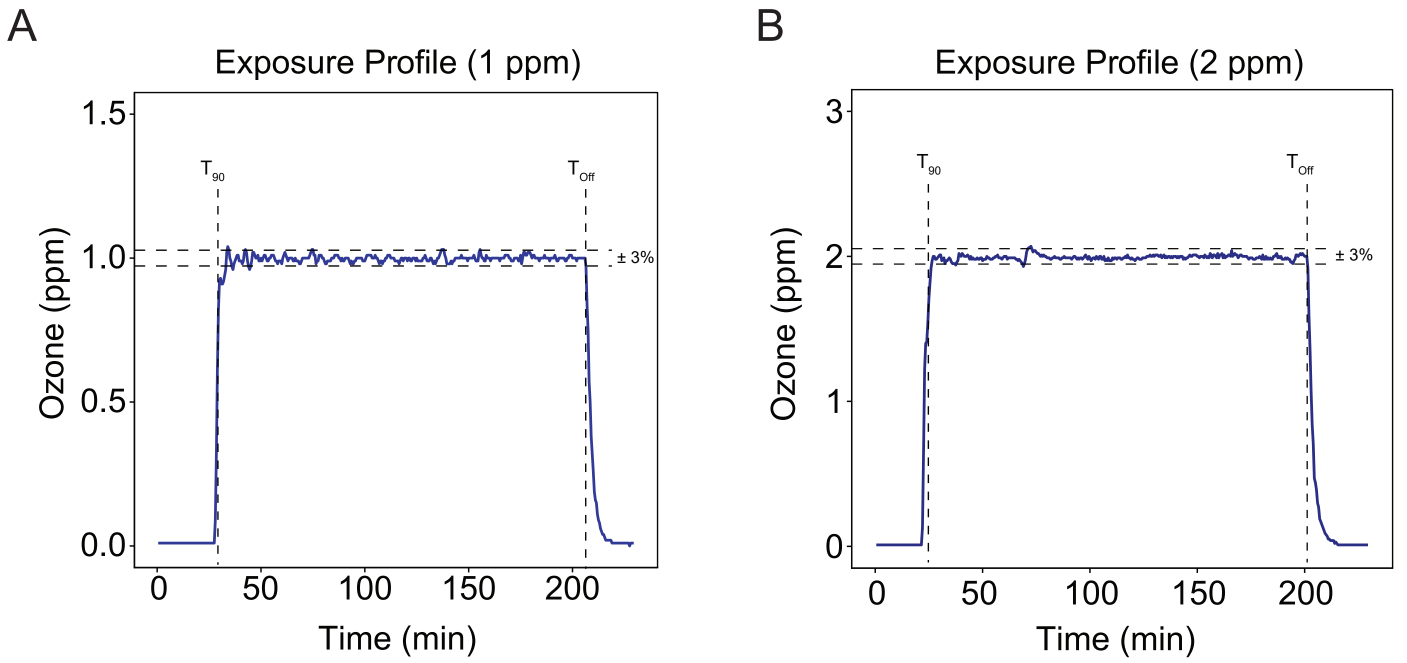

**Supplemental Figure 2.**

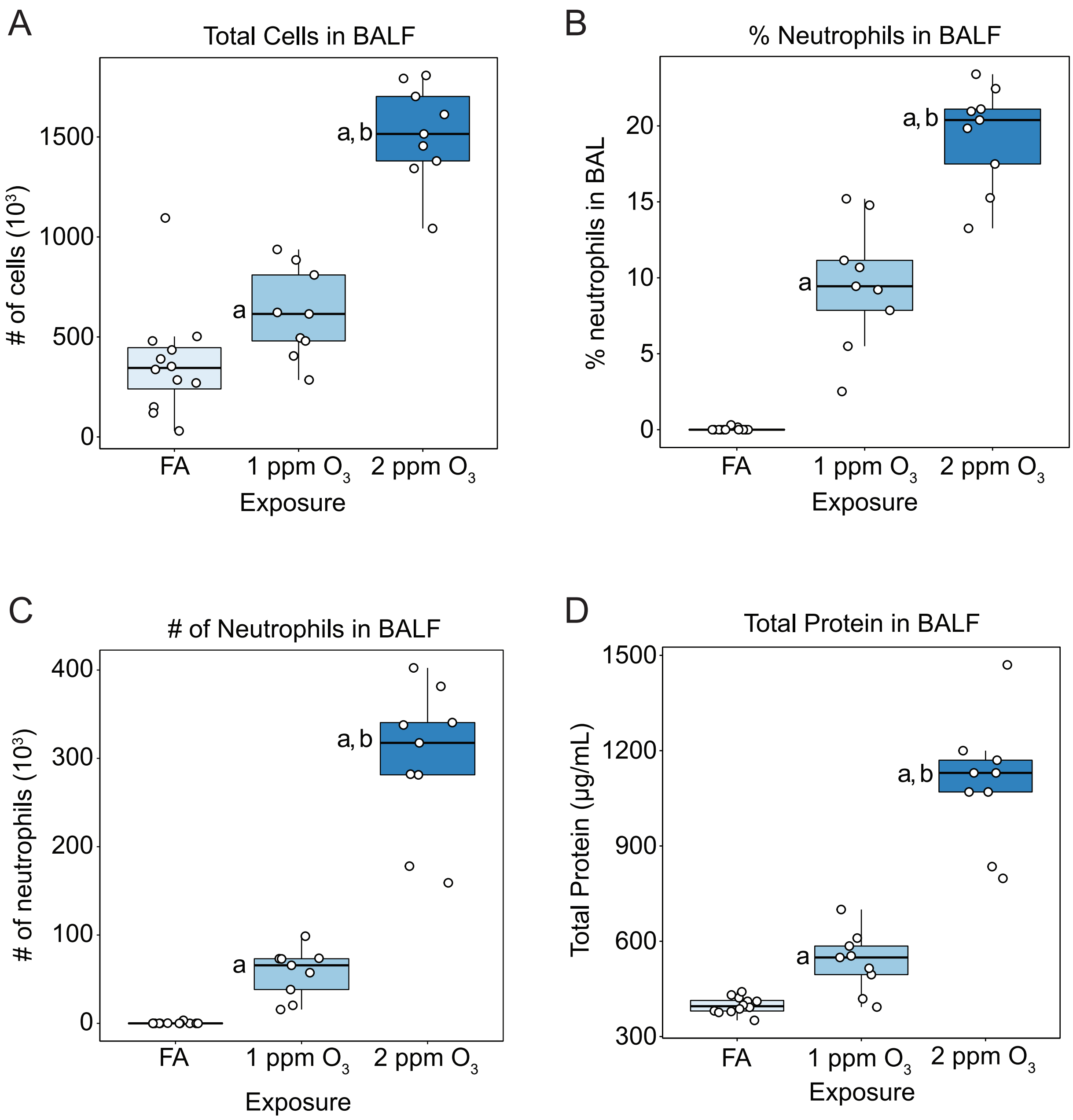

**Supplemental Figure 3.**

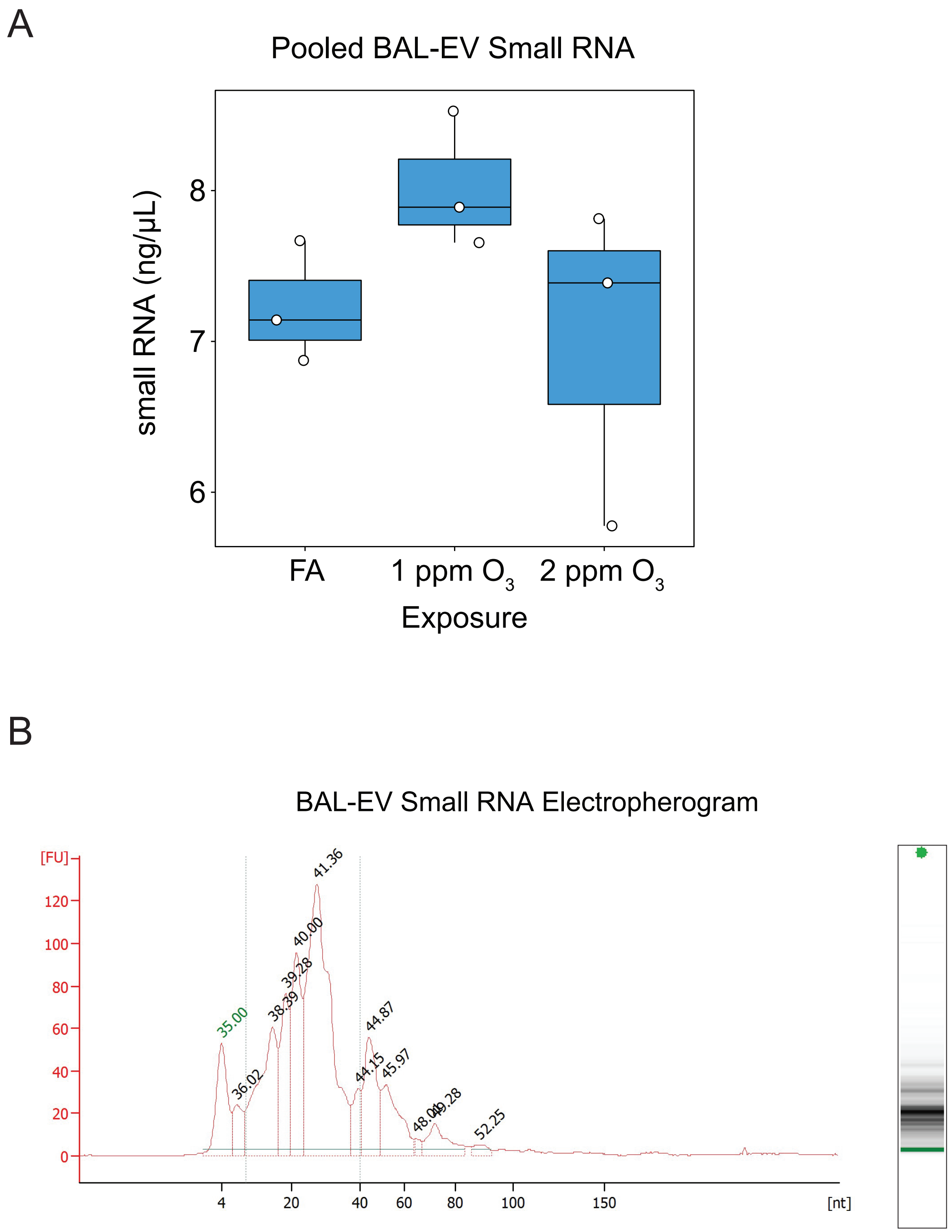

**Supplemental Figure 4.**

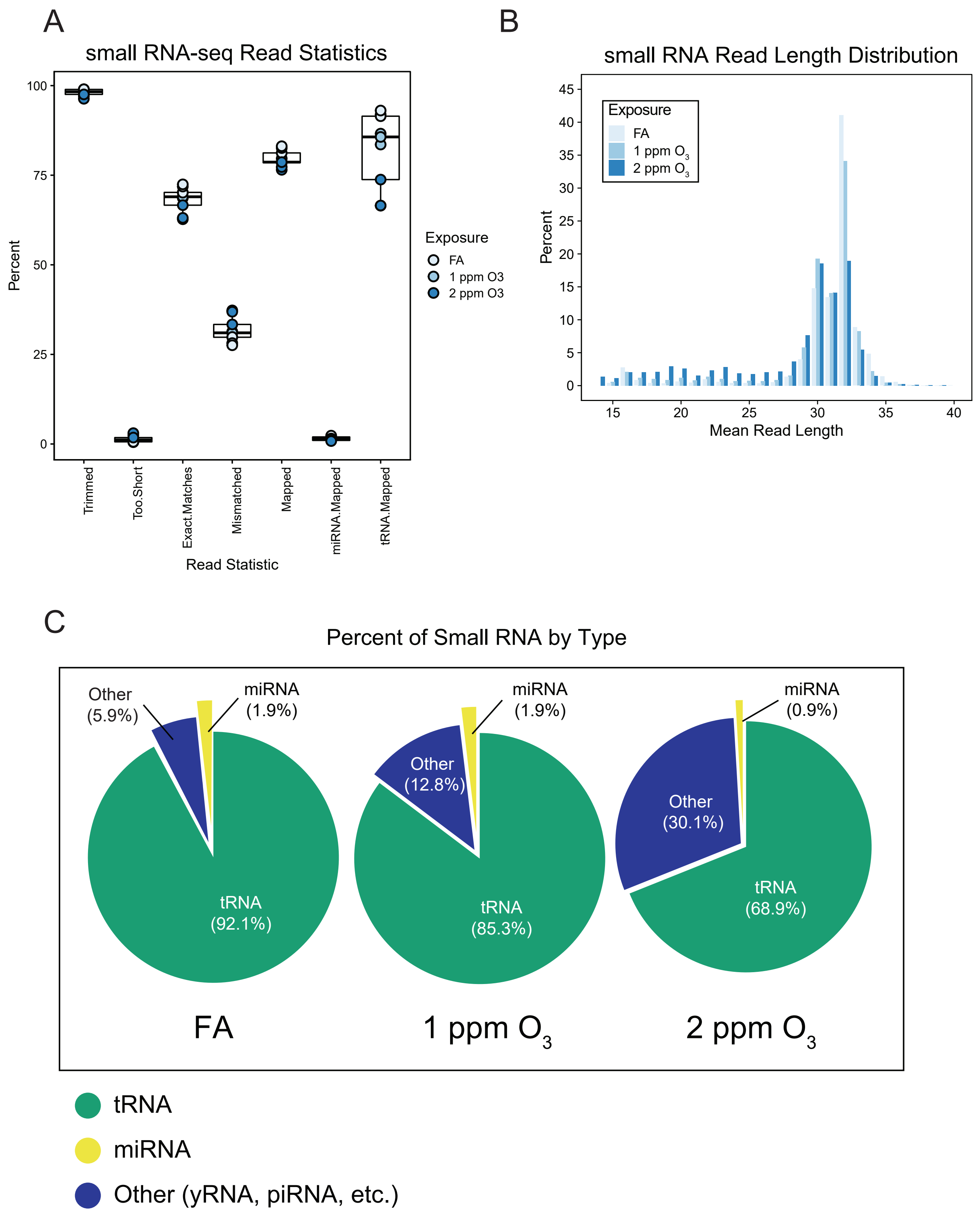

**Supplemental Figure 5.**

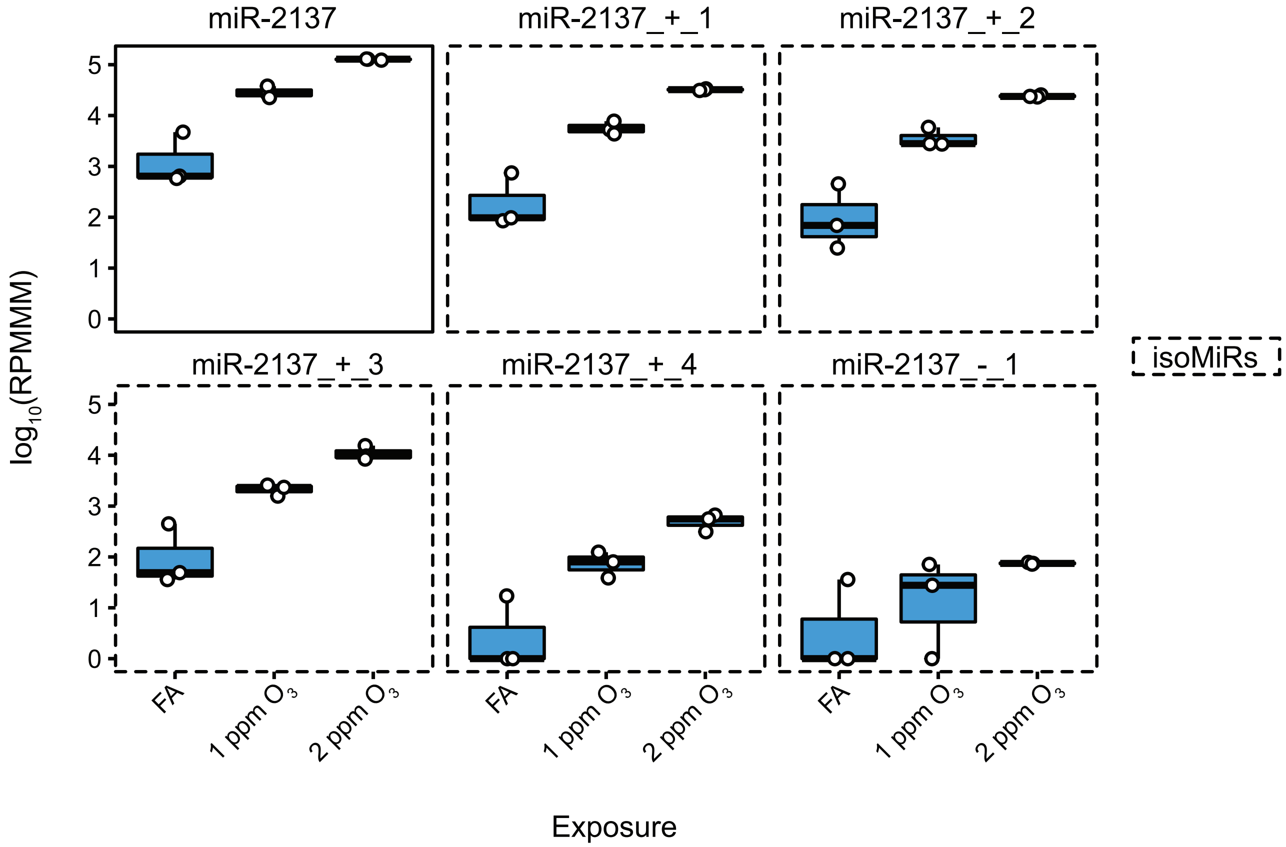

**Supplemental Figure 6.**

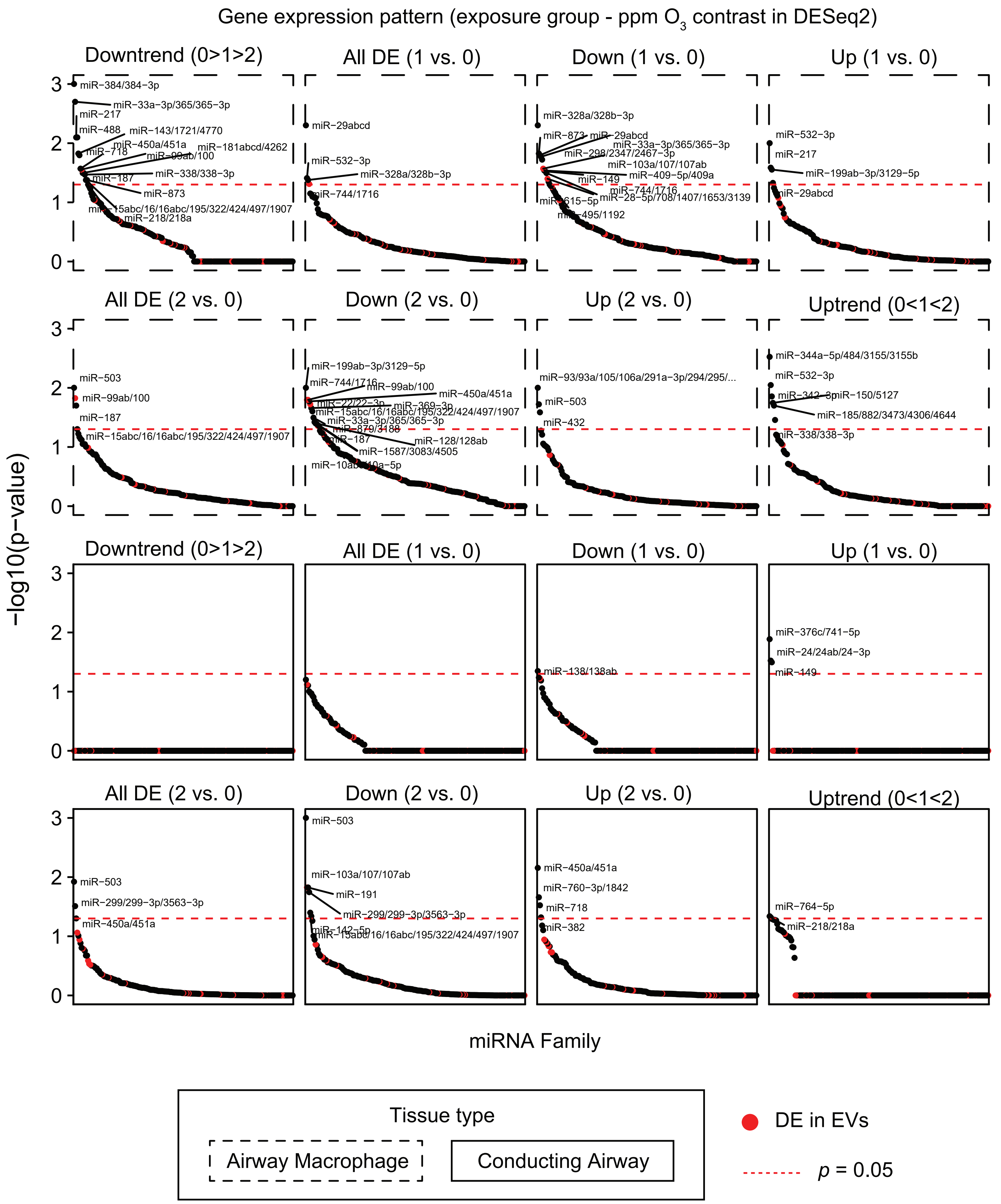

**Supplemental Figure 7.**

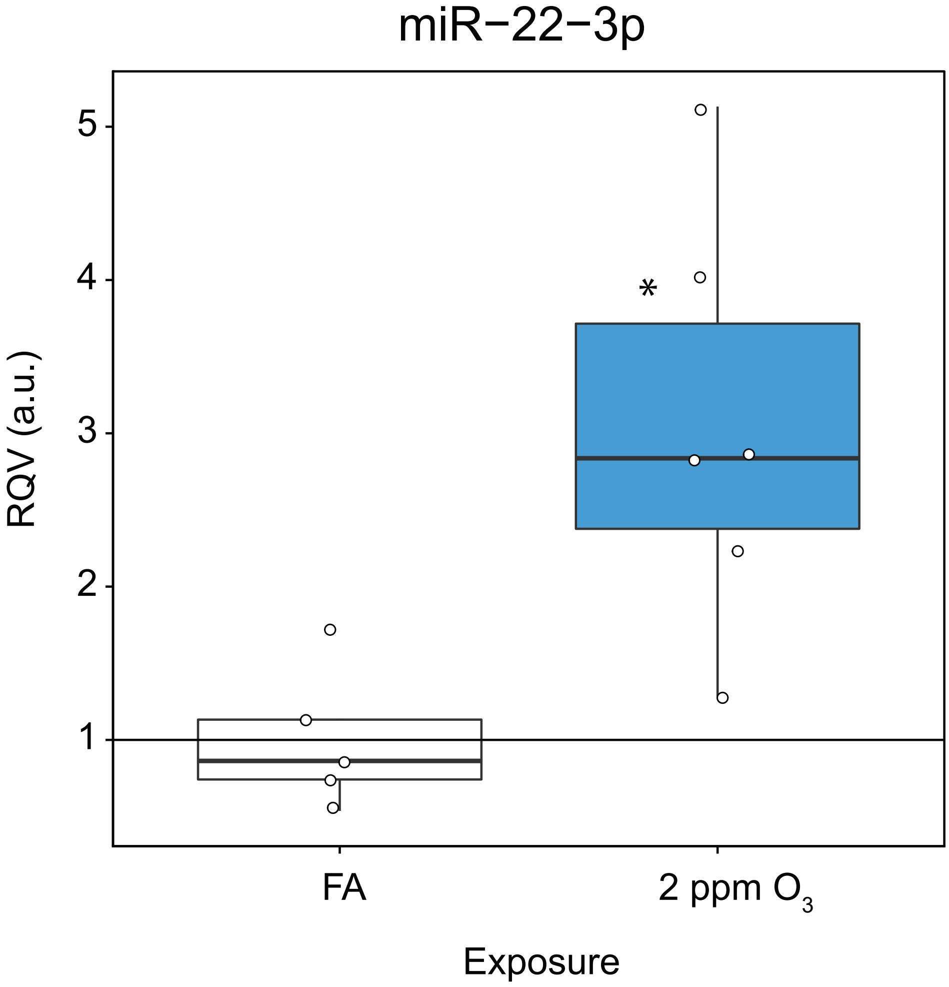
